# Empirical Analyses of Associations between State Political Attitudes and Gains in Health Insurance Coverage after the Affordable Care Act (ACA)

**DOI:** 10.1101/605022

**Authors:** Bisakha Sen, Reena Joseph

## Abstract

**Objectives:** To explore whether state-level political-sentiment is associated with gains in insurance post Affordable Care Act (ACA). This is especially relevant given the lawsuit brought by several Republican-leaning states against the ACA, and the ruling of one Texas federal judge that the ACA is unconstitutional, which potentially jeopardizes ACA’s future.

**Methods:** Multivariate linear-probability models are estimated using data from the Behavioral Risk Factor Surveillance Systems for 2011-2017. The outcome is self-reported insurance status. States are placed in quartiles based on votes for President Obama in 2008 and 2012 elections. Starting 2014, ACA health exchanges became active and several states expanded Medicaid, so 2014 onwards is considered as the ‘post-ACA’ period. Models are estimated for all adults under 65-years and for young adults under 35-years. All models control extensively for respondent socio-economic-demographic characteristics and state characteristics.

**Results:** In the pre-ACA baseline period, respondents in states with higher Anti-Obama-voting (AOV) were less likely to have insurance. For example, residents in highest AOV-quartile states were 8.0-percentage-points less likely (p<0.001) to have insurance than those in the lowest AOV-quartile states. Post-ACA, fewer high AOV-quartile states expanded Medicaid, and overall insurance gains inclusive of Medicaid-expansion are similar across states. However, net of Medicaid-expansion, residents in higher AOV states saw higher insurance gains. For example, all adults had 2.8-percentage points higher likelihood (p<0.01) and young adults had 4.9-percentage point higher likelihood (p<0.01) of getting insurance in the highest AOV-quartile states compared to the lowest AOV quartile states. Minorities and those with chronic-conditions had larger insurance gains across the country post-ACA, but the extent of these gains did not differ by state AOV levels.

**Conclusions:** State AOV and insurance gains from ACA appear to be incongruent. Policymakers and stakeholders should be aware that non-Medicaid residents of higher-AOV states might potentially lose the most if ACA is revoked.

## Introduction

The Patient Protection and Affordable Care Act (hereafter ACA) very substantially changed the health insurance landscape in the United States, and scientific research indicates that although there were significant increases in rates of health insurance coverage in the U.S. after its passage, yet opinions about the ACA remain deeply divided along political lines [1–6]. The Kaiser Health Tracking Poll [7] shows that overall unfavorable ratings of ACA dropped from a high of 53% in July 2014 to 40% in November 2018. However, among those who identify as Republicans, approximately 75% continue to view it unfavorably, while less than 20% of those who identify as Democrats and about 45% of those who identify as Independents do so. The ACA’s future also remains uncertain under the current administration. One key provision, the ‘individual mandate’ that requires individuals to purchase insurance or pay a fine, has already been revoked. Essential health Benefits (EHB) in the ACA continue to be targeted by Republicans, and may be particularly vulnerable in states where health exchanges are struggling with insurance companies pulling out [8]. The current administration declined to defend the ACA against a lawsuit [9] brought by several states that claim that the provision requiring insurance companies to cover individuals with pre-existing conditions and at no higher premiums is unconstitutional – and on December 14, 2018, a federal judge in Texas ruled that the ACA was unconstitutional in the absence of the individual mandate. The case is very likely to go to the Supreme Court, and the future of the ACA is likely to be a key issue in the upcoming Presidential elections.

It is no secret that the political landscape in the U.S. has become increasingly partisan over the past decades. The ACA is President Obama’s signature policy and strongly identified with him, as reflected in its moniker ‘Obamacare’. Public opinion regarding the ACA, whether it should be repealed and whether the federal government should have a role in ensuring health coverage for all, are all strongly patterned along political affiliation [10, 11]. It is also no surprise that the 20 states that brought the aforementioned lawsuit are largely Republican leaning. In this context, it is of particular interest to see if the gains in insurance coverage after key provisions of ACA went into effect vary by a state’s political leanings.

Previous studies have found that after ACA health exchanges went into effect and some states expanded Medicaid in 2014, nationwide uninsurance rates dropped substantially, and that gains in insurance were divided equally between gains in Medicaid and private coverage [2–4, 6, 12, 13]. One study has also documented substantial variation across states in gains following ACA [3]. However, to our knowledge, no study has explicitly looked at whether the across-state variation in coverage gain is associated with political opinions in that state. In this study, we address this gap. We operationalize the states’ political leanings by measuring the extent of what can be termed ‘Anti-Obama voting’ (AOV) in each state, and explore whether AOV is associated with relative gains in insurance coverage following ACA. We are particularly interested in gains in insurance net of state Medicaid expansion, since the latter was largely predicted by the political affiliations of the state legislatures and governors. We argue that gains in insurance-coverage after controlling for Medicaid expansion better captures voluntary take-up of insurance by state residents through the ACA health exchanges, and that it is more pertinent to explore if and how such voluntary take-up is influenced by state AOV.

We also look at whether the relative gains by those with pre-existing conditions compared to their healthier counterparts, and gains by minorities compared to non-Hispanic whites, vary by state AOV. Existing research indicates that minorities gained more under ACA than non-Hispanic whites [14–16]. Since minorities are more likely to identify as Democrats than non-Hispanic whites [17], it may be that in states with high AOV, voluntary take-up of insurance was disproportionately driven minorities, or by those with pre-existing conditions who deem health insurance as more of a necessity than their healthier counterparts. The findings will help inform on which states’ residents may be more impacted should the ACA be revoked or effectively crippled.

## Methods

We use 2011-2017 data from the Behavioral Risk Factor Surveillance System (BRFSS) for our analyses. BRFSS is an annual telephone survey conducted by state health departments and the US Centers for Disease Control and Prevention (CDC) that collects data on socio-demographic characteristics, health insurance status, preventive services, risky behaviors, and self-assessed health for all 50 states and the District of Columbia. A random digit dialing method is used to select a representative sample of respondents from the noninstitutionalized adult population. BRFSS has been extensively used to research health impacts of the ACA [5, 18–21]. It is particularly appropriate because its large number of observations, more than 300,000 per year, allows us to decipher the effects of the ACA. This is important since only a fraction of the population is directly affected by the legislation. We confine our sample to 19- to 64-year-olds since the ACA was not designed to impact senior Medicare recipients. We also exclude pre-2011 years because from 2011 the BRFSS included cell phones in its sampling. As individuals who exclusively use cell phones are disproportionately young, this inclusion essentially changes the characteristics of the sample, as well as the sample means of several key variables pertaining to health and health insurance status. We omit residents of Massachusetts from our analyses, since Massachusetts passed its own health reform law in 2006 under Governor Mitt Romney, providing almost universal coverage to its residents. We also omit the District of Columbia.

### Measures

We use multivariate linear probability regression models with individual-level data pooled over time and state. Outcome and control variables are described below, and descriptive statistics are shown in Table 1.

**Table 1:** Descriptive statistics for outcome, individual characteristics and state controls, by AOV quartile.

| Variables | AOV_Q1<br>(N=430,615) | AOV_Q2<br>(N=560,979) | AOV_Q3<br>(N=552,121) | AOV_Q4<br>(N=529,766) |
| --- | --- | --- | --- | --- |
| **Individual level variables** |  |  |  |  |
| Health Insurance Coverage | 0.891 | 0.881 | 0.830 | 0.852 |
| Health Insurance Coverage pre-ACA | 0.864<br>(N=183,311) | 0.851<br>(N=255,069) | 0.801<br>(N=246,527) | 0.822<br>(N=242,937) |
| Health Insurance Coverage post-ACA | 0.911<br>(N=246,264) | 0.906<br>(N=304,155) | 0.854<br>(N=303,678) | 0.877<br>(N=285,175) |
| Diagnosed with Chronic condition | 0.413 | 0.433 | 0.440 | 0.447 |
| Self-reported not good health | 0.145 | 0.145 | 0.181 | 0.176 |
| Age-group (in years) |  |  |  |  |
| 25-34 | 0.150 | 0.146 | 0.148 | 0.153 |
| 35-44 | 0.184 | 0.179 | 0.176 | 0.183 |
| 45-54 | 0.261 | 0.254 | 0.250 | 0.246 |
| 55-64 | 0.326 | 0.342 | 0.345 | 0.335 |
| Black | 0.098 | 0.049 | 0.127 | 0.077 |
| Hispanic | 0.130 | 0.081 | 0.091 | 0.050 |
| Other race | 0.137 | 0.078 | 0.081 | 0.065 |
| Female | 0.561 | 0.552 | 0.569 | 0.571 |
| Low income | 0.277 | 0.293 | 0.346 | 0.313 |
| Missing Family Income | 0.130 | 0.119 | 0.132 | 0.136 |
| Salaried employment | 0.585 | 0.582 | 0.544 | 0.560 |
| Education level |  |  |  |  |
| Grades 1-8 (Elementary) | 0.025 | 0.014 | 0.023 | 0.018 |
| Grades 9-11 (Some high school) | 0.044 | 0.041 | 0.062 | 0.055 |
| Grades 12 or GED (High school graduate) | 0.242 | 0.257 | 0.283 | 0.293 |
| College 1 year to 3 years (Some college or technical school) | 0.252 | 0.289 | 0.286 | 0.301 |
| College 4 years or more (College graduate) | 0.431 | 0.394 | 0.340 | 0.329 |
| Refused | 0.005 | 0.003 | 0.003 | 0.003 |
| Marital Status |  |  |  |  |
| Divorced | 0.130 | 0.134 | 0.146 | 0.142 |
| Widowed | 0.033 | 0.031 | 0.041 | 0.038 |
| Separated | 0.028 | 0.021 | 0.031 | 0.024 |
| Never married | 0.234 | 0.201 | 0.200 | 0.175 |
| A member of an unmarried couple | 0.044 | 0.042 | 0.035 | 0.028 |
| Refused | 0.008 | 0.007 | 0.007 | 0.005 |
| **State level variables** |  |  |  |  |
| State poverty rates | 11.402<br>(2.069) | 12.670<br>(2.200) | 15.787<br>(1.948) | 14.473<br>(2.945) |
| State unemployment rates | 6.478<br>(2.265) | 6.115<br>(2.177) | 6.304<br>(2.234) | 5.545<br>(1.829) |
| Percent Hispanic | 16.027<br>(10.089) | 11.071<br>(11.362) | 12.395<br>(12.112) | 7.811<br>(3.514) |
| Percent Black | 12.680<br>(8.647) | 6.348<br>(5.270) | 14.314<br>(10.251) | 8.411<br>(8.754) |
| Percent Urban | 86.010<br>(13.837) | 72.279<br>(12.017) | 70.684<br>(13.509) | 67.122<br>(10.638) |
| Percent with Bachelors or higher | 32.516<br>(2.422) | 29.174<br>(3.971) | 24.968<br>(1.763) | 24.820<br>(3.716) |
| Democrat Governor | 0.92<br>(0.451) | 0.458<br>(0.498) | 0.191<br>(0.393) | 0.184<br>(0.387) |
| Number of Democrat Senators | 1.771 | 1.693 | 0.739 | 0.444 |
|  | (0.420) | (0.610) | (0.651) | (0.656) |
| State expanded Medicaid | 1 | 0.858 | 0.388 | 0.268 |
| State exchange or state-federal partnership | 0.763 | 0.712 | 0 | 0.224 |

#### Outcome

The primary outcome of interest in all cases is self-reported insurance status at the time of the survey (insured versus uninsured). BRFSSS asks whether the respondent has health insurance, but in most years does not ask the source of coverage. Thus, we cannot decipher whether health insurance was purchased from a health exchange. However, our interest is in overall changes in health insurance coverage, since it could be argued that gains in insurance via health exchanges might be offset by reductions in employer-provided insurance.

#### Post-ACA

The primary independent variable is a binary indicator for ‘Post-ACA’, which is 1 for 2014-2017, and 0 before, since the exchanges went into operation in 2014.

#### Anti-Obama Voting

We operationalize AOV by taking the simple average of the percentage of the state’s popular vote that were cast for a presidential candidate other than Barack Obama in 2008 and 2012. We also do sensitivity analyses using just the percentage of votes in 2008, since it might be argued that votes in 2012 were endogenous to how people felt about the ACA. We identify the four quartiles in the distribution of votes cast, and categorize states by which quartile they fall into. These are hereafter referred to as AOV-Q1 (first quartile), AOV-Q2 (second quartile), AOV-Q3 (third quartile) and AOV-Q4 (fourth quartile). Thus, states in the fourth or highest quartile can be considered as having the highest levels of AOV, and states in the first or lowest quartile having the lowest level of AOV (or, being most ‘pro’ President Obama). The states in each quartile are listed in Appendix A.

#### Chronic Pre-existing Conditions

The BRFSS asks if the respondent has been diagnosed with any of an extensive list of chronic conditions. We create a binary indicator that captures if they responded in the affirmative to at least one of the following diagnosis: diabetes, heart attack, coronary heart disease, stroke, asthma, cancer, chronic obstructive pulmonary disease, arthritis, kidney disease or depressive disorders.

#### Other individual characteristics

We use the standard demographic controls at the individual level, including age, race-ethnicity, gender, marital status, education, income level, and salaried employment.

#### Time-trend

The economy was improving and expanding over 2011-2017, which could likely lead to greater insurance coverage from other sources like employer-provided insurance; hence we add a time-trend variable to account for coverage growth from such sources.

#### Other state characteristics

We present models with and without controls for whether the state expanded Medicaid in 2014. The latter models show overall gain in insurance after 2014. However, we are primarily interested in the former models, which arguably better indicate the gains due to voluntary take-up of insurance through health exchanges. We also control for if the exchange was state-operated or a state-federal partnership, as opposed to the reference group of federal exchanges. To help reduce bias from a state’s socio-economic-demographic characteristics, we control for the percentage of the population who are African-American, the percent who are Hispanic, percentage in poverty, percent living in an urban area, the percent with a college degree in 2011, and the percent unemployed. To account for other facets of the state’s political climate, we control for whether the state’s governor over this period was a Democrat and the party affiliation of the two Senators from the state in different years.

Descriptive statistics for all of the variables are in Table 1.

### Regression Models

We use multivariate linear probability regression models and a variation of the conventional ‘difference-in-difference’ approach. Detailed equations are in the appendix. The key results of interest pertain to coefficient estimates of the interactions between the binary ‘Post-ACA’ variable and the state AOV quartiles (AOV-Q_s_). AOV-Q_1_ – the most ‘pro-Obama’ quartile of states – is the reference group. Essentially, this approach allows us to test whether there were statistical differences in insurance gains between the AOV-Q_1_ states and states in each of the other AOV-Q_s_ groups after 2014. Results are presented in Table 2.

**Table 2:** Linear probability models for health insurance changes after ACA, across different state AOV levels.

| Outcome: Has Health Insurance | Full Sample<sup>(A)</sup><br>(N=1484884) | Under 35 years<sup>(A)</sup><br>(N=329499) | Full Sample<br>(N=1484884) | Under 35 years<br>(N=329499) |
| --- | --- | --- | --- | --- |
| AOV-Q1 | Ref group | Ref group | Ref group | Ref group |
| AOV-Q2 | -0.0557**<br>(-21.53) | -0.0650**<br>(-12.51) | -0.0534**<br>(-22.11) | -0.0633**<br>(-13.03) |
| AOV-Q3 | -0.0911**<br>(-23.98) | -0.107**<br>(-13.82) | -0.0791**<br>(-25.77) | -0.0931**<br>(-15.18) |
| AOV-Q4 | -0.0854**<br>(-20.55) | -0.103**<br>(-12.32) | -0.0802**<br>(-22.85) | -0.100**<br>(-14.29) |
| post-ACA<sup>(B)</sup> | 0.0055<br>(1.25) | -0.0154<br>(-1.71) | 0.0453**<br>(15.16) | 0.0456**<br>(7.80) |
| AOV-Q2*post-ACA | 0.0086**<br>(2.79) | 0.0248**<br>(4.03) | -0.00353<br>(-1.32) | 0.00194<br>(0.37) |
| AOV-Q3*post-ACA | 0.0314** | 0.0483** | -0.00199 | -0.00263 |
|  | (7.78) | (5.88) | (-0.66) | (-0.45) |
| AOV-Q4*post-ACA | 0.0282** | 0.0488** | 0.00491 | 0.0116* |
|  | (7.28) | (6.23) | (1.60) | (1.91) |
| *N* | 1484884 | 329499 | 2065195 | 470977 |

*T*-statistics in parentheses. All models include controls for the full list of individual demographic and socio-demographic characteristics as well as all the state level characteristics that are listed in Table
\* $p < 0.10$ , \* $p < 0.05$ , \*\* $p < 0.01$
(A): Models also include controls for Medicaid expansion and whether the exchange was a state-run exchange.
(B): These results reflect changes post-ACA for the reference state category AOV-Q<sub>1</sub>

Next, we investigate whether minorities and those with chronic conditions gained differentially compared to their counterparts in states with different AOV levels. For this, we use a variation of the ‘triple-difference’ approach in the econometric literature. Again, the detailed equations are in the appendix. Essentially, this method allows us to test whether the relative gains in insurance by a particular sub-group (minority or chronic condition) in each of the higher AOV-Q_s_ states are statistically different than the relative gains for that sub-group in the AOV-Q_1_ (i.e. reference group) states.

We also run models to see if minorities and those with chronic conditions gain relatively more in insurance coverage in the country as a whole. This is primarily to compare our findings with studies with fewer post-ACA years of data, and the results are given in the appendix.

Models are estimated for the full sample of 19-64 years olds, and also just for those under 35 years old (hereafter referred to as young adults), since the latter traditionally has less access to employer-sponsored health insurance, and have been shown to overall be more impacted by ACA. The statistical software Stata (v.14) is used, and the SVY routine applied to account for sampling design in BRFSS. α is set at 0.05 for purpose of hypothesis testing.

## Results

Appendix A lists the states in each AOV quartile. Respectively, mean votes against President Obama in AOV-Q_1_ states was 37.8 percent (range: 28.8 percent-42.2 percent), in AOV-Q_2_ states were 46.6 percent (range: 43 percent – 48.1 percent), in AOV-Q_3_ states were 54.3 percent (range: 49 percent – 58.4 percent), and in AOV-Q_4_ states were 62.9 percent (range: 59.6 percent – 70.3 percent).

Descriptive statistics in Table 1 show that residents in AOV-Q_1_ and AOV-Q_2_ states have higher rates of overall health insurance, but that they also had higher rates of insurance in the pre-ACA years. For example, 86.4% and 85% of respondents in AOV-Q_1_ and AOV-Q_2_ states respectively reported having health insurance pre-ACA, whereas 80.1% and 82.2% in AOV-Q_3_ and AOV-Q_4_ states report the same. Residents in all four quartiles of states show an approximately 5-percentage point increase in the likelihood of insurance post-ACA compared to pre-ACA.

Unsurprisingly, more AOV-Q_1_ and AOV-Q_2_ residents saw Medicaid expansion in their states (100% and 85.8% respectively) as well as state health exchanges (76.3% and 71.2% respectively). In comparison, just 38.8% of AOV-Q_3_ residents and 26.8% AOV-Q_4_ residents saw Medicaid expansion in their states; 0% AOV-Q_3_ state residents and 22.4% AOV-Q_4_ state residents had state exchanges.

Some individual-level differences by AOV quartile also stand out. For example, in AOV-Q_1_ states, about 27.8% respondents have family incomes under $35,000 annually, and 41.3% have ever been diagnosed with a chronic condition; whereas in AOV-Q_4_ states, 31.3% are low-income, and 44.7% have a chronic condition diagnosis and 17.6% self-report not good health. The discrepancy in income level may partly explain the lower rates of pre-ACA insurance. On the other hand, no clear patterns in the relationship between AOV quartile and minority status emerge -- with AOV-Q_1_ and AOV-Q_3_ states having more minority respondents compared AOV-Q_2_ and AOV-Q_4_ states.

Table 2 shows regression results for gains in the likelihood of insurance after ACA for residents in the different AOV-Q groups. Pre-ACA, compared to residents of AOV-Q_1_ states, AOV-Q_2_ state residents were approximately 5.6 percentage points less likely to be insured (p<0.01), AOV-Q_3_ state residents were approximately 9 percentage points less likely to be insured (p<0.01) and AOV-Q_4_ state residents were approximately 8.5 percentage points less likely to be insured (p<0.01). These pre-ACA differences are slightly larger for the young adults. When Medicaid expansion is not controlled for, residents of AOV-Q_1_ states show an increase in insurance coverage post-ACA (β= 0.045, p<0.01) and there are no statistical differences in the gains by their counterparts from other AOV-Q groups. However, after controlling for Medicaid expansion, there is no significant increase in insurance coverage likelihood for AOV-Q_1_ state residents (β=0.005, p>0.10), but there are significant additional gains for residents of AOV-Q_2_ (β= 0.008, p<0.01), AOV-Q_3_ (β= 0.031, p<0.01) and AOV-Q_4_ states (β=0.028, p<0.01). The gains are even higher in models that just consider young adults (β=0.025, p<0.01; β=0.048, p<0.01, and β=0.050, p<0.01 respectively). This strongly suggests that in AOV-Q1 states, insurance gains were largely attributable to Medicaid expansion. However, in higher AOV states, larger gains likely happened from other sources -- such as take-up of insurance through the ACA health exchanges.

We confirm (results are in appendix) findings from previous studies [13–16] that nationwide, blacks and Hispanics were less likely to have insurance coverage in the pre-ACA period than non-Hispanic whites were. However, compared to non-Hispanic whites, they also made larger gains in coverage post-ACA. Those with chronic conditions were more likely to have insurance compared to their healthy counterparts pre-ACA; nonetheless, they also showed larger increases in insurance coverage post –ACA.

Table 3 presents key results from the quasi triple-difference model when Medicaid expansion is controlled for (full results are in the appendix). In the AOV-Q1 reference group, respondents with chronic conditions have a 1.1 percentage point greater likelihood (p<0.05) than their healthier counterparts of gaining coverage. However, there is no evidence of statistically greater relative gains for the other AOV-Q_s_ groups. In fact, the relative gains appear to be lower in AOV-Q_3_ states, though it is not a statistically significant difference (β= −0.011, p>0.10). Black respondents have a 2.6 percentage point (4.5 percentage points for young adults) greater likelihood of gaining coverage than non-Hispanic whites in AOV-Q_1_ states, and this is not statistically different from relative gains by Black respondents in other AOV-Q_s_ states. Similarly, Hispanics have a 2 percentage point greater likelihood of gaining insurance (p<0.05) compared to non-Hispanic whites in AOV-Q1 states in the full sample – though no relatively higher gains are seen for young adult Hispanics. Again, there is no statistical difference in relative gains by Hispanics in other AOV-Q_s_ states. Thus, those with chronic conditions, and racial and ethnic minorities, make greater relative gains in insurance coverage compared to their counterparts after ACA (this is also corroborated by the countrywide results in the appendix), but the extent of those relative gains do not differ significantly by state AOV levels.

**Table 3:** Quasi triple difference models for changes in insurance for respondents with chronic conditions, and minority respondents, across different state AOV levels.

| Outcome: Has Health Insurance | Full Sample<sup>(A)</sup><br>(N=1,484,884) | Under 35 years<sup>(A)</sup><br>(N=329,499) |
| --- | --- | --- |
| Variation Across AOV quartile states for changes in coverage for those with chronic conditions |  |  |
| post-ACA*chronic <sup>(B)</sup> | 0.0113* | 0.0194* |
|  | (2.16) | (1.78) |
| AOV-Q2*post-ACA*chronic | 0.0005 | -0.0025 |
|  | (0.08) | (-0.19) |
| AOV-Q3*post-ACA*chronic | -0.0110 | -0.0202 |
|  | (-1.52) | (-1.34) |
| AOV-Q4*post-ACA*chronic | 0.0087 | 0.0162 |
|  | (1.23) | (1.07) |
| Variation Across AOV quartile states for changes in coverage by race and by ethnicity |  |  |
| post-ACA*Black <sup>(B)</sup> | 0.0262** | 0.0447** |
|  | (3.20) | (2.76) |
| AOV-Q2*post-ACA*Black | 0.0115 | -0.00116 |
|  | (0.97) | (-0.05) |
| AOV-Q3*post-ACA*Black | 0.0063 | -0.0113 |
|  | (0.57) | (-0.52) |
| AOV-Q4*post-ACA*Black | -0.0089 | -0.0344 |
|  | (-0.74) | (-1.47) |
| post-ACA*Hispanic <sup>(B)</sup> | 0.0193* | 0.0004 |
|  | (2.54) | (0.03) |
| AOV-Q2*post-ACA*Hispanic | 0.0104 | 0.0164 |
|  | (0.90) | (0.88) |
| AOV-Q3*post-ACA*Hispanic | 0.00886 | 0.0128 |
|  | (0.78) | (0.69) |
| AOV-Q4*post-ACA*Hispanic | -0.0105 | 0.0117 |
|  | (-0.71) | (0.51) |

Only the results for how health insurance changed for the sub-group of interest compared to counterparts, and across different state AOV levels are shown. Full tables are in Appendix.
*T*-statistics in parentheses. All models include controls for the full list of individual demographic and socio-demographic characteristics as well as all the state level characteristics that are listed in Table
\* $p < 0.10$ , \* $p < 0.05$ , \*\* $p < 0.01$
<sup>(A)</sup>: Models also include controls for Medicaid expansion and whether the exchange was a state-run exchange.
<sup>(B)</sup>: These results show the relative change post-ACA for the sub-group of interest compared to their counterparts in the reference state category AOV-Q

We repeated all of the above analyses by re-defining AOV-Q_s_ only using votes against Obama in the 2008 presidential elections – since it could be argued that his support in 2012 was partly driven by attitudes about ACA. Our results remain virtually identical to those presented with that alternate specification.

## Discussion and conclusions

The ACA provides multiple avenues for expanding health insurance. Starting September 2010, young adults could stay on parental plans via dependent coverage up to age 26. Health exchanges started providing coverage starting January 2014, and included protections against differential treatment for those with pre-existing conditions, premium subsidies on a sliding scale up to 400% of the federal poverty level (FPL), and individual mandates to incentivize healthy individuals to purchase insurance. States that elected to expand Medicaid covered all adults up to 133% of Federal Poverty Limit (FPL). That these provisions expanded insurance coverage is evidenced by several studies using datasets such as Gallup-Healthways Wellbeing Index [2, 12], the National Health Interview Survey [14], the American Community Survey [3], the Commonwealth Fund Survey [22] and BRFSS [5, 13, 23].

There is also evidence of heterogeneity in gains. Early provisions like extending parental insurance coverage to young adults benefitted young, white adults more than ethnic minorities [23]. However, after the health exchanges started operating and some states expanded Medicaid, racial and ethnic minorities and low-income adults gained the most in coverage [3, 14–16]. One study looked at state by state gains immediately after 2014 and found the highest gains were in Kentucky and Nevada and the lowest was in Wyoming [4]. Several studies have looked at gains for states that expanded versus did not expand Medicaid coverage, as well as variation in gains across income groups [2, 4, 22], and at least one study has documented lower cost burdens on low-income patients when Medicaid has been expanded [24].

No existing study has explicitly looked at whether there are variations in gains by political sentiment in the state. This is pertinent for many reasons. Even in 2016, attitudes about ACA were strongly patterned on political affiliation, with the Gallup Poll reporting that 82 percent of Republican or Republican-leaning respondents opposed keeping the ACA in place [25]. In 2018, twenty states with Republican legislatures and/or governors filed a lawsuit to declare unconstitutional the pre-exiting conditions protections in ACA, and were partially victorious given the ruling by the Texas federal judge. The survival of ACA and the protections it offers is likely to be a leading issue in the 2020 Presidential elections. This leads to the question of which states’ residents gained the most after the ACA was implemented, and, by extension, would be hurt the most if it was revoked.

We explored whether gains in insurance coverage after ACA were associated with the state’s political leanings -- which we operationalized as support or lack thereof for President Obama in the presidential elections as measured by share of votes. There are several reasons to speculate why insurance gains may be associated with AOV. For example, individuals in higher AOV states may have negative opinions about President Obama and any programs initiated by him, may be more likely to hear elected state officials disparage the ACA, or may have less access to healthcare navigators tasked with informing people about provisions in the ACA [26, 27]. High AOV states were mostly less likely to expand Medicaid, so there would be fewer avenues for adults below the federal poverty level to gain coverage in those states. On the other hand, if the higher AOV states also had more people with relatively poor access to insurance prior to the ACA and hence greater unmet need for coverage –– then more people in these states might sign up for insurance via exchanges if the ACA provided such opportunities.

Our results seem to support the latter conjecture. Individuals in higher AOV states were significantly less likely to have health insurance compared to those in the lowest AOV quartile states before the health exchanges started operating in 2014. For example, even after accounting for individual socio-demographic factors, income and work status, SES, the average adult in AOV-Q4 states was 8.0 percentage points less likely to have health insurance, and the average young adult (that is, <35 years) was about 10 percentage points less likely to have health insurance, than their counterparts in AOV-Q_1_ states. After 2014, gains across states were similar when including Medicaid expansion. However, net of Medicaid expansion, the AOV-Q_1_ states actually showed the least gains. In contrast, there were statistically significant higher gains among states in the higher AOV quartiles. The largest gains were for young adults in AOV-Q_3_ and AOV-Q states, whose likelihood of coverage increased by about 5 percentage points compared to the reference group. This tentatively suggests that there was more unmet need for health insurance in higher AOV states, and relatively greater number people in these states signed up for health insurance through the exchanges.

One might speculate that these higher gains in insurance in AOV-Q_3_ and AOV-Q_4_ states may be disproportionately driven by people with pre-existing conditions, who consider insurance a necessity and are hence relatively impervious to political opinions on ACA. Alternatively, one might speculate that the higher gains may be driven by racial and ethnic minorities, who tend to lean Democrat politically [17] and also may have less access to employer-provided insurance. However, our quasi-triple difference analyses indicate that while the relative gains for these sub-groups are higher overall, they are not statistically different across state AOV quartiles. Thus, the greater gains in insurance in high AOV states cannot be attributed to disproportionately larger gains for sicker residents or minority residents.

### Limitations

We acknowledge several limitations. BRFSS does not inform on the source of health insurance coverage for its full sample – so we cannot decipher whether or not insurance was obtained through an ACA health exchange. We deliberately include in our models several controls to help control for employer-provided health insurance – such as the individual’s status as a salaried employee, the state unemployment rate, and a time-trend to account for economic growth and recovery from the Great Recession over the study period. This strengthens our conjecture that, once Medicaid expansion is also controlled for, the remaining changes in insurance that we see post-2014 are likely driven by participation in health exchanges; however, we cannot ascertain it beyond doubt. One other inherent challenge in studying the effects of ACA is that there is no obvious ‘control group’ of ineligible people who are otherwise reasonably similar to those eligible to participate in ACA, which also limits our ability to make causal inferences about whether observed changes are caused purely by ACA. We acknowledge the usual problems with self-reported data. We also acknowledge that our definition of pre-existing conditions as those reporting one of several chronic conditions in the survey may be problematic, since the diagnosis itself may be linked to having a medical check-up in the past – thus chronic conditions may be underreported for those without insurance. We further note that political sentiment is a complex and multi-faceted issue that can potentially be measured many other ways, and results may be sensitive to how it is operationalized. We use the share of votes against President Obama averaged over 2008-2012, and also do sensitivity analyses using just the 2008 votes, but we also recognize that a sizeable share of the population does not vote and hence this measure does not accurately capture their political leanings. BRFSS itself does not provide any information on political opinions and affiliations of individual respondents, so we cannot use that information to validate our findings. Finally, we focus health insurance in this paper, and cannot comment on access, quality of health services provided, or use of preventive care, nor can we decipher whether there were long-term improvements in the health of respondents due to the improved insurance coverage.

In conclusion, this study strongly suggests there is likely a disconnect between political opinions in a state and the actual gains in insurance coverage for the state’s residents. Arguably, since lack of health insurance coverage was an issue that only affected approximately 15 percent of the U.S. population prior to the ACA, the issue of uninsurance might not be of high importance to the average voter when deciding their political affiliation. That said, one could argue that state legislators, regardless of party affiliation, should be more concerned about the possible impact on their more vulnerable constituents if the ACA was revoked or effectively crippled. Concerns have already been raised about how repealing the ACA would adversely affect the opioid epidemic in the U.S [28]. The majority of the U.S. population currently support protecting health insurance coverage for people with chronic conditions [29], and in the most recent midterm elections, four Republican-leaning states – Idaho, Montana, Nebraska and Utah – voted on Medicaid expansion ballot initiatives, and it passed in all states except Montana. Yet, the political environment remains volatile and the December 14, 2018 ruling by the Texas federal judge puts the future of the ACA in renewed jeopardy. This study’s findings underline the imperative need to inform policymakers on the impact of ACA in states across the political spectrum, and the repercussions if it is revoked, which should help inform the debate about the future of healthcare in the U.S.

